# Earliest evidence for frugivory and seed dispersal by birds

**DOI:** 10.1101/2022.03.01.482456

**Authors:** Han Hu, Yan Wang, Paul G. McDonald, Stephen Wroe, Jingmai K. O’Connor, Alexander Bjarnason, Joseph J. Bevitt, Xuwei Yin, Xiaoting Zheng, Zhonghe Zhou, Roger B.J. Benson

**Author notes:** For correspondence (Y.W.); (H.H.).

## Abstract

The Early Cretaceous diversification of birds was a major event in the history of terrestrial ecosystems, occurring during the earliest phase of the Cretaceous Terrestrial Revolution. Frugivorous birds play an important role in seed dispersal today, and may have done so since their origins. However, evidence of this has been lacking. *Jeholornis* is one of the earliest-diverging birds, only slightly more derived than *Archaeopteryx*, but its cranial anatomy has been poorly understood, obscuring diet-related functional interpretations. Originally hypothesised to be granivorous based on seeds preserved as gut contents, this interpretation has become controversial. We conducted high-resolution synchrotron tomography on an exquisitely preserved new skull of *Jeholornis*, revealing remarkable cranial plesiomorphies combined with a specialised rostrum. We use this to provide a near-complete cranial reconstruction of *Jeholornis*, and exclude the possibility that *Jeholornis* was granivorous, based on morphometric analyses of the mandible (3D) and cranium (2D), and comparisons with the 3D alimentary contents of extant birds. We show that *Jeholornis* was at least seasonally frugivorous, providing the earliest evidence for fruit consumption in birds, and indicating that seed dispersal was present from early in the avian radiation. As highly-mobile seed dispersers, early frugivorous birds could expand the scope for biotic dispersal in plants, and may explain, in part, the subsequent evolutionary expansion of fruits, indicating a potential role of bird-plant interactions in the Cretaceous Terrestrial Revolution.

## Introduction

Birds are among the most speciose extant vertebrate groups, playing unique ecological roles through their diverse flight and dietary adaptations (Prum et al., 2015). Modern birds include both specialised and opportunistic frugivores, that collectively are major consumers of fruits and important agents of seed dispersal. The early ecological diversification of birds in the Early Cretaceous (>130 Ma) (Yang et al., 2020) was a landmark event in the evolution of terrestrial ecosystems, adding considerably to species richness of terrestrial ecosystems (Benson, 2018; Yu et al., 2021), and with impacts on the evolutionary histories of other flying groups (Benson et al., 2014b; Clapham and Karr, 2012). The origin and early diversification of birds in the Early Cretaceous was followed by a considerable long-term expansion of the abundance and disparity of fruits and fruit-like structures (Eriksson et al., 2000a; Eriksson, 2008), as part of the major floral transition from gymnosperm-to

Due to its key phylogenetic position, *Jeholornis* has been frequently studied and cited, and many specimens are known. However, because specimens are often compressed, and are preserved in slabs, little unequivocal cranial information has been available (Lefèvre et al.,

2007; Rauhut et al., 2018). The palatine is broad with a well-developed jugal process that contacts the maxilla. The pterygoid is elongated with no sign of the shortening that occurs in more derived birds and the pterygoid flange is well-developed, indicating the presence of an ectopterygoid. The vomer is dorsoventrally thin with bifurcated caudal flanges oriented nearly vertical to the rostral body, similar to the condition in *Sapeornis* (Hu et al., 2019).

While the temporal and palatal regions retain plesiomorphies, the rostrum of *Jeholornis* is heavily modified. The new specimen reveals that its premaxillae corpora are fused while the frontal processes remain separate. Rostral fusion of the premaxillae is also present in extant birds, confuciusomithiforms and several enantiomithines e.g., *Linyiornis* and *Shangyang* (Wang and Zhou, 2019; Wang et al., 2016). Its occurrence in *Jeholornis* indicates that it occurred phylogenetically earlier than previously thought. *Jeholornis* also shows dental reduction, with an edentulous premaxilla, two rostrally restricted maxillary teeth and three

process seeds using a gastric mill, with minimal beak processing; (4) Fruit eaters; and (5) Other diets (such as folivores, carnivores and omnivores).

### Mandibular Morphospace

The principal components analysis (PCA) results reveal that a large portion of mandibular shape variation (PC1: 36%) is related to the relative length of the mandible compared to its rostral depth: positive values of PC1 indicate short, deep mandibles, whereas negative values indicate long, low mandibles. PC2 explains 34% of variation and is also related to the relative depth of the mandible, with positive values indicating low mandibles with coronoid eminence absent or less developed, and negative values indicating deep mandibles with a large coronoid eminence. PC3 (10% of variation) is related to the curvature, with positive values indicating a straight profile in lateral view, and negative values indicating rostroventral curvature of the rostral portion of the mandible (Figure 2A, B).

The results plot *Jeholornis* near the centre of mandibular morphospace. Seed-crackers, especially parrots, are clearly separated from the other diet types including *Jeholornis* in mandibular morphospace. They occupy a distinct region with high, positive values of PC1 and low, negative values of PC2, reflecting their deep and anteroposteriorly short mandibles with a large coronoid process and deep mandibular symphysis, which suits their seed-cracking diet by reducing the beak failure risk during cracking (Soons et al., 2015, 2010). The

is related to their ability to de-husk seeds (Meij and Bout, 2008). Therefore, finches are also clearly distinct from the position of *Jeholornis* in mandibular morphospace (Figure 2A, B), rejecting the previous hypothesis of *Jeholornis* as a seed cracker (both parrot-type and finch-type) (Zhou and Zhang, 2002).

*Jeholornis* is plotted within the overlapping range of frugivores, seed-grinders, and birds

crackers have relatively short, deep and rostroventrally curved rostra compared to most other birds, including *Jeholornis, Sapeornis* and other Mesozoic taxa.

Mesozoic taxa are mostly separated from modern birds along PC2 and PC3, occupying negative values of PC2 and positive values of PC3 separately (Figure 2C, D). Among them, *Jeholornis* and *Sapeornis* are more similar to extant birds along PC2, which describes rostral morphology. This may reflect the dietary specialization of *Jeholornis* and *Sapeornis* (frugivory-like or granivory) compared to other Mesozoic taxa. Nevertheless, they cluster with other Mesozoic taxa along cranial PC3, indicating conservative aspects shared with nonavian theropods, especially a proportionally small orbit and external naris.

### Alimentary Content Analyses

remains are highly fragmented in seed-cracking parrots (Figure 4E), whereas in seed-cracking passerines, although the crop contents are almost intact, those in the stomach are also highly fragmentary (Figure 4D, Figure 4 - figure supplements 1E, F). This is consistent with behavioural observations of finches and other granivorous passerines (Billerman et al., 2020), in which seed-cracking passerines use the beak only to remove the outer coats of seeds, and do not fragment the seed before ingestion, differing from parrots that can fragment seeds prior to ingestion (Figure 4E). Fragmentation (in granivorous passerines) is completed later by the gastric mill, showing a similar condition of seed-grinders. Fragmentation of seeds in passerines is primarily achieved through the gastric mill, similar to some seed-grinders e.g. *Ectopistes migratorius* (Passenger pigeon) (Figure 4C, Figure 4 - figure supplements 1B). However, in most seed-grinders the gut contents consist of abraded and partially-damaged, rather than highly-fragmented, seed remains (Figure 4B, Figure 4 - figure supplements 2A-F).

Seed remains in all the sampled granivores were tightly aggregated together, and typically co-occurred with gastroliths (Figure 4B-E). Gastroliths are especially abundant in some seed-grinders and seed-cracking passerines (Figure 4B, D) compared to the parrot (Figure 4E) and pigeon (Figure 4C). In contrast, the seed remains in frugivores are

Digital reconstruction of an exceptionally well-preserved new specimen of the early-diverging bird *Jeholornis* reveals a plesiomorphic, diapsid skull, sharing numerous features with non-avian theropods. These features include a complete postorbital bar, unreduced squamosal, and unmodified palate (Hu et al., 2020b, 2020a, 2019; Rauhut et al., 2018), reinforcing evidence for an early-diverging phylogenetic position among birds (Wang et al., 2018; Zhou and Zhang, 2002). Nevertheless, compared to *Archaeopteryx* (Rauhut, 2014; Rauhut et al., 2018), *Jeholornis* also possesses clear diet-related specialisations of the rostrum including partial fusion of the premaxillae and a strongly reduced dentition.

Our GMM analyses reveal that the mandibular and cranial shapes of *Jeholornis* and *Sapeornis* are distinct from those of seed-cracking granivorous birds, consistent with earlier assumptions that the delicate, vestigial dentary teeth of *Jeholornis* would be too prone to

diversification, with birds enabling seed dispersal for plants, and obtaining a rich energy resource in return. New discoveries and comparative analyses are required to further test this hypothesis, by insights into the ecologies of more early bird species, and the potential role of the birds during the transition from gymnosperm-to angiosperm-dominated floras.

## Materials and Methods

### Taxonomy of *Jeholornis* STM 3-8

*Jeholornis* STM 3-8 was collected from the Jiufotang Formation (~120 Ma) (He et al., 2004) at the Dapingfang locality in Chaoyang, Liaoning province, preserving a complete and mostly articulated skull, and a few postcranial elements. This new specimen is tentatively assigned to *Jeholornis prima* based on the presence of the following features: relatively robust mandible with three rostrally restricted teeth; edentulous and robust premaxilla; maxilla lacking teeth in the caudal portion; long bony tail consisting of more than 20 caudal vertebrates. This specimen could be distinguished from *Jeholornis palmapenis* by its flattened dorsal margin of ilium, compared to the strongly convex condition in *J. palmapenis*

Microtomographic measurements of *Jeholornis* STM 3-8 were performed using the Imaging and Medical Beamline (IMBL) at the Australian Nuclear Science and Technology

the location of the preorbital ossifications being the highest uncertainty. The reassembled cranial model was then used as the reference for the 2D reconstruction of the *Jeholornis* skull in lateral and ventral views (Figure 1E, F).

**Figure 1.**
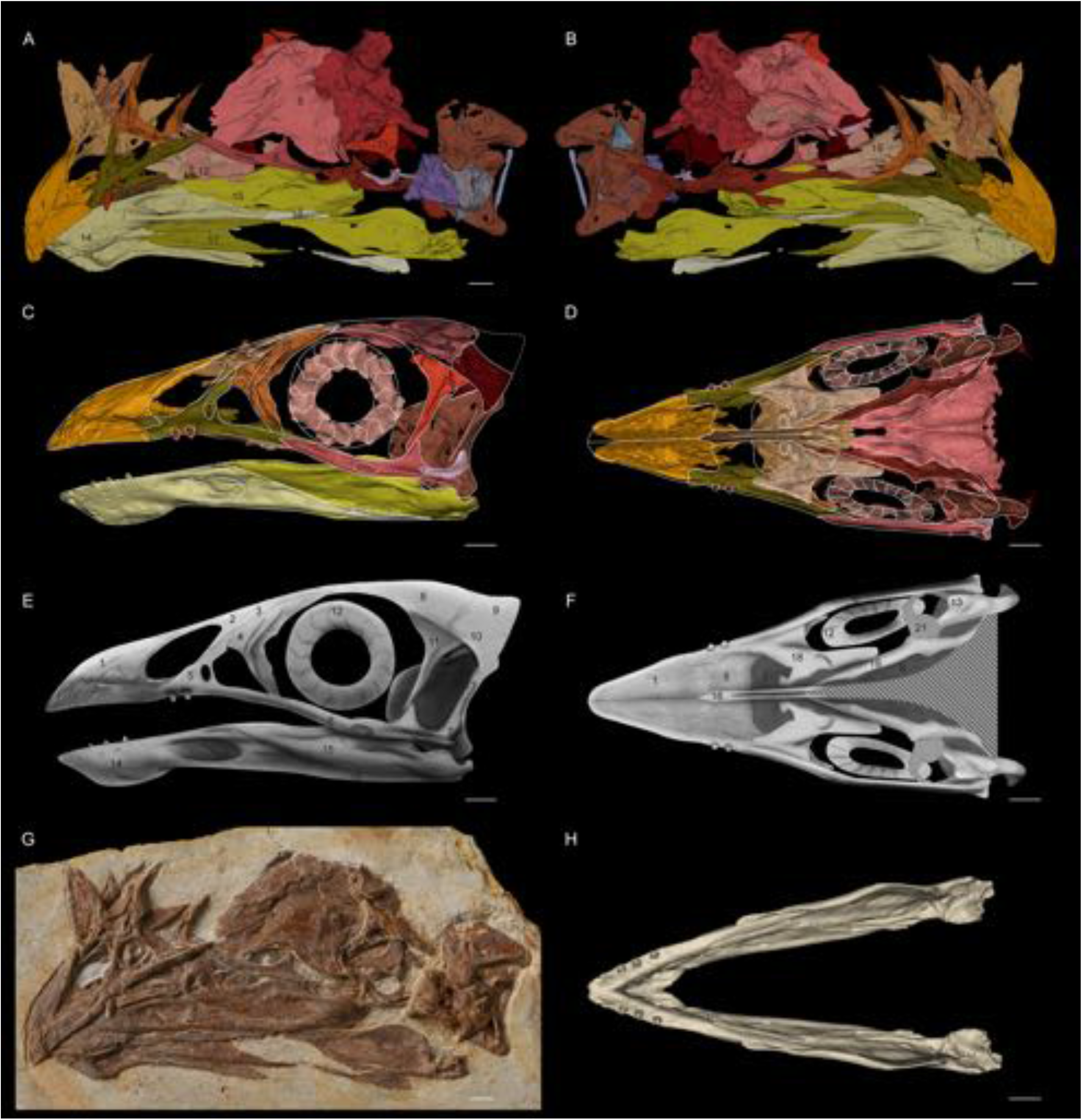
*Jeholornis* STM 3-8. **(A)** Left and **(B)** right view of the 3D reconstructed model of the skull. **(C)** Left and **(D)** ventral view of the reassembled 3D model of the skull. **(E)** Left and **(F)** ventral view of the 2D cranial reconstruction. **(G)** Photograph of the skull. **(H)** Dorsal view of the reassembled 3D model of the mandible. Abbreviations: 1. premaxilla; 2. nasal; 3. preorbital ossification; 4. lacrimal; 5. maxilla; 6. jugal; 7. quadratojugal; 8. frontal; 9. braincase; 10. squamosal; 11. postorbital; 12. scleral ring; 13. quadrate; 14. dentary; 15. surangular; 16. angular; 17. splenial; 18. vomer; 19. palatine; 20. pterygoid; 21. potential ectopterygoid. Different bones are indicated by different colours. Scale bar equals 5mm.

**Figure 2.**
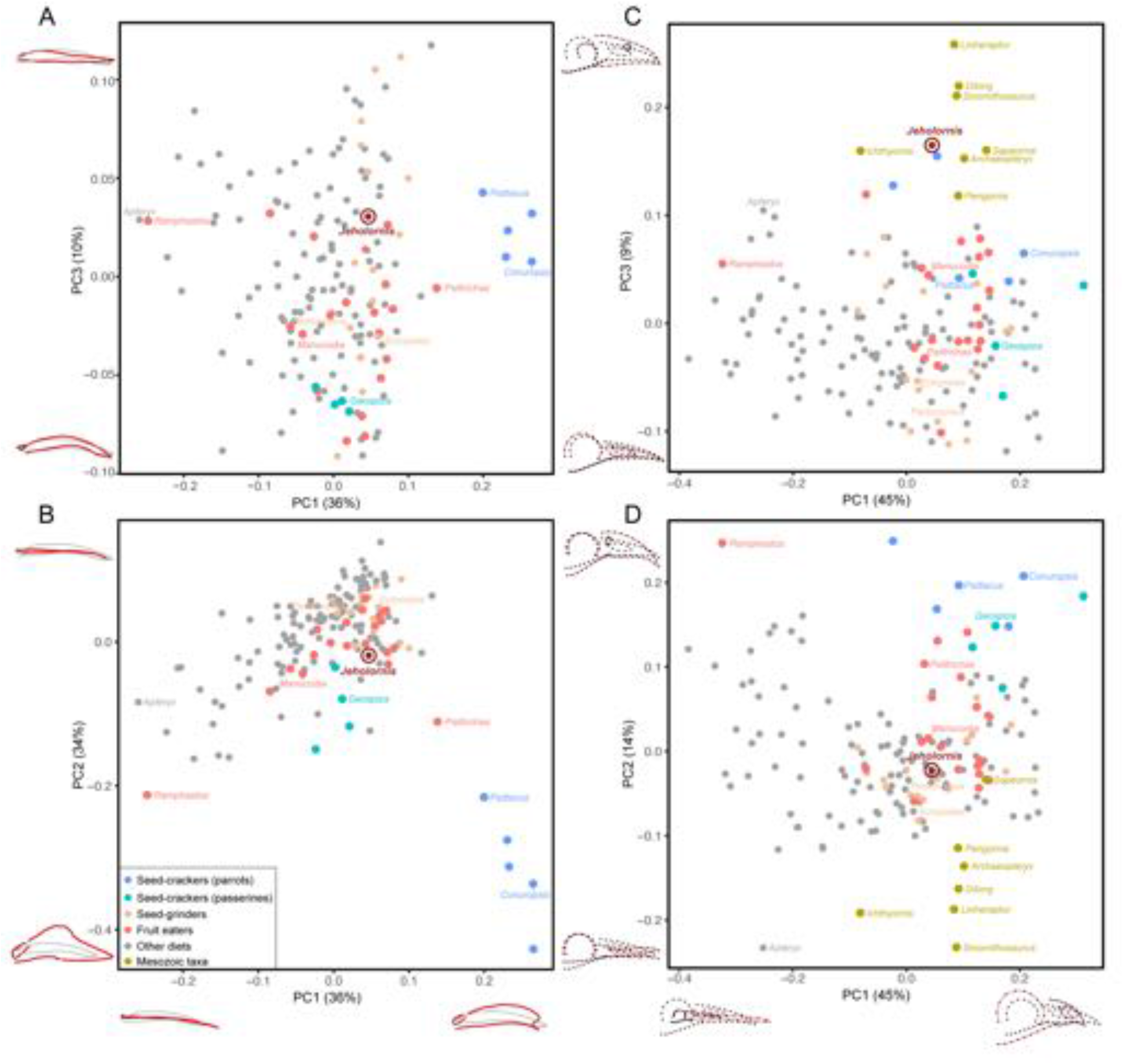
PCA result of 3D mandible shape (A, B) and 2D skull shape (C, D). Different diet categories are indicated by different colours, and key samples are labelled with generic names.

**Figure 3.**
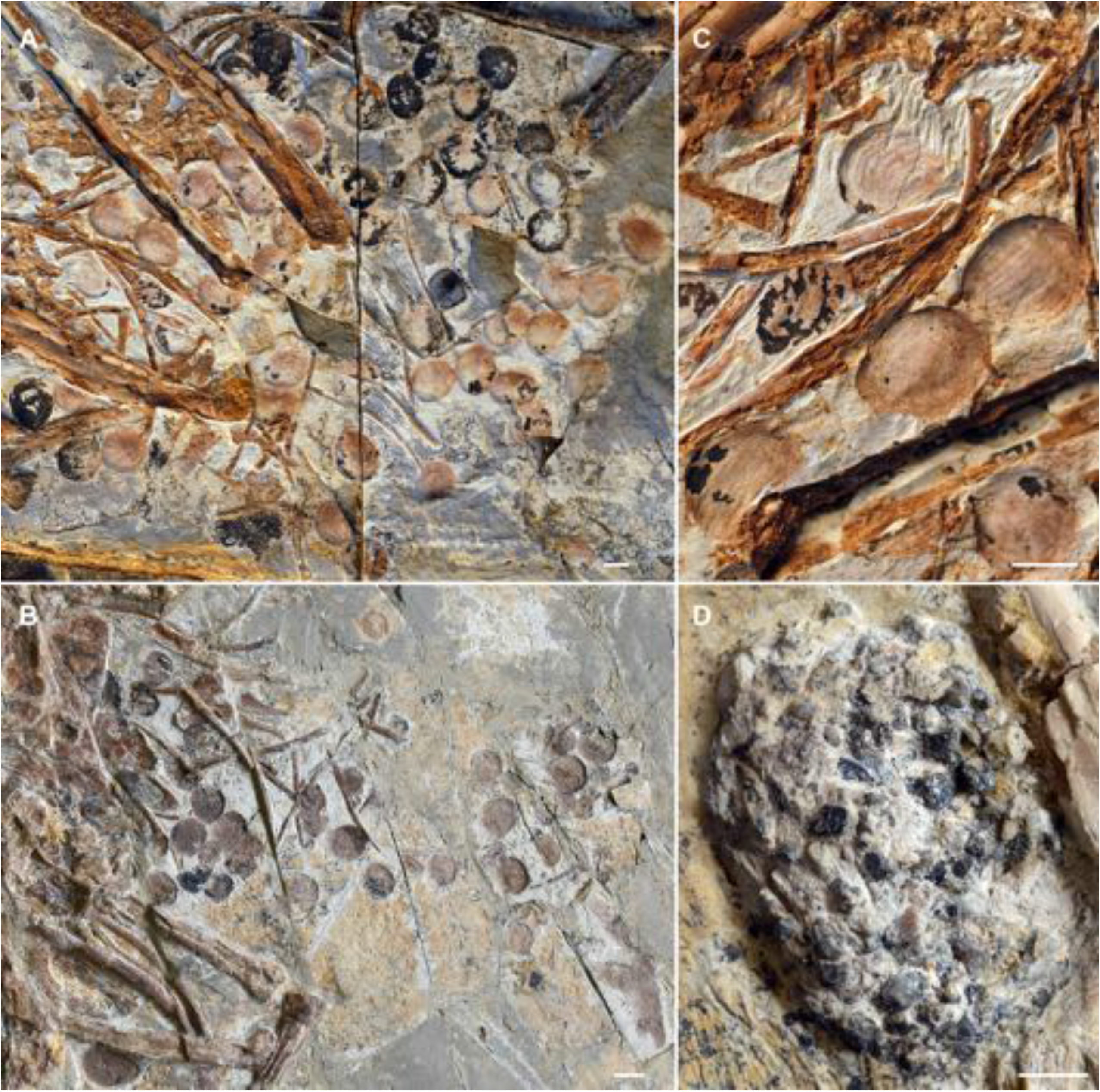
Seeds preserved in the abdominal area of selected *J. prima* specimens. **(A)** IVPP V13274 (holotype). **(B)** STM 2-41. **(C)** Close up image of seeds in IVPP V13274 (A). **(D)** Gastrolith mass in *J. Prima* STM 2-15. Photos in A-D followed Figures in reconstructed models. Scale bars equal 5mm for the whole models and slices, and 1mm for the the magnification boxes.

**Figure 4.**
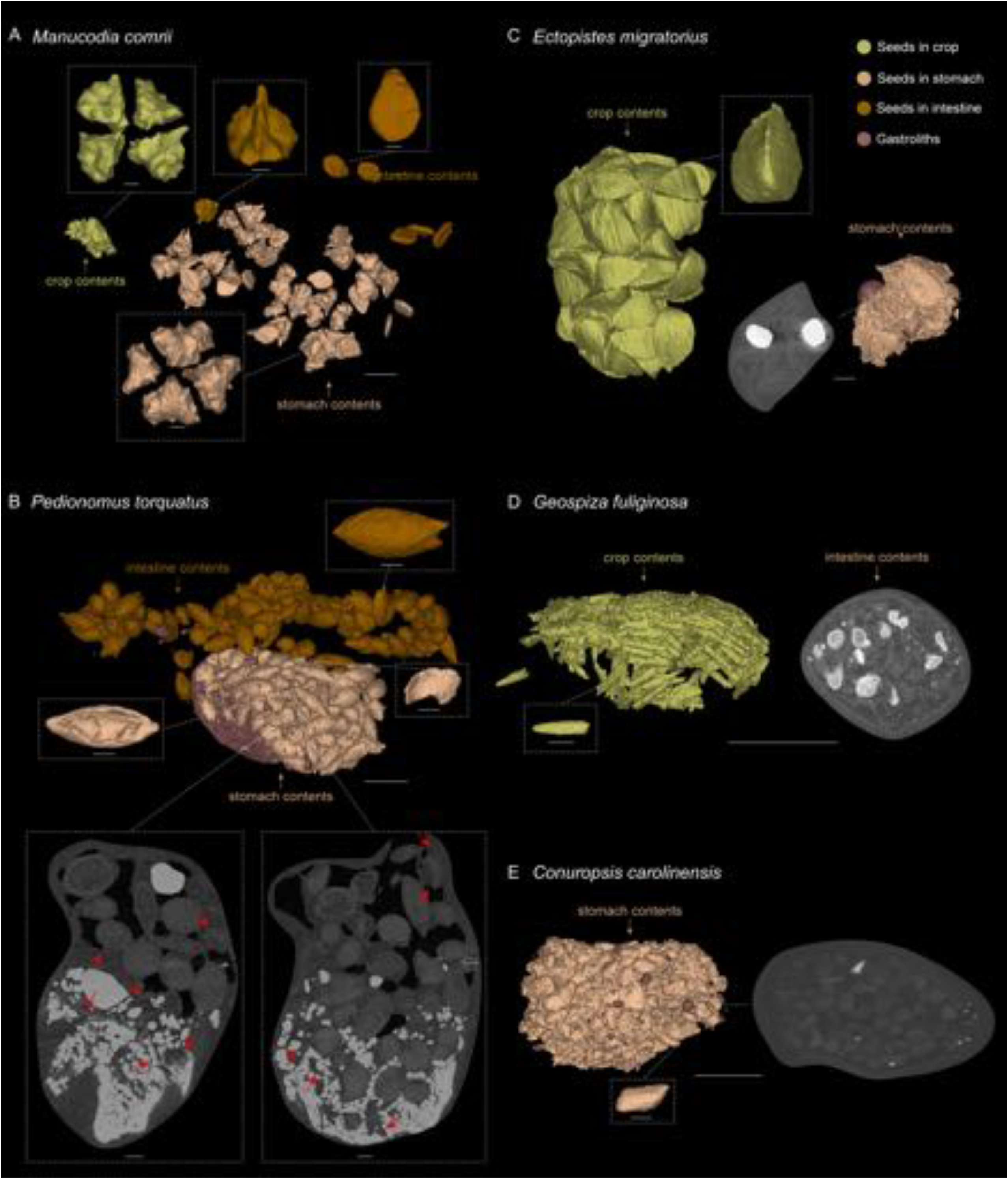

### GMM analyses

The dataset incorporates *Jeholornis* and 160 extant bird species representing 111 families and 36 orders in our 3D mandible analysis, with additional Mesozoic theropods in 2D skull analysis including: *Sinornithosaurus* (Dromaeosauridae) (Xu and Wu, 2001), *Linheraptor* (Dromaeosauridae) (Xu et al., 2015), *Dilong* (Tyrannosauroidea) (Xu et al., 2004)’ *Archaeopteryx* (non-Ornithothoraces Aves) (Rauhut, 2014), *Sapeornis* (non-Omithothoraces Aves) (Hu et al., 2019), *Pengornis* (Enantiomithes)

1. Frugivores: *Manucodia comrii* (Curl-crested manucode, Figure 4A, Figure 4 - figure supplements 1A), is a specialized fruit eater (Billerman et al., 2020). Several whole fruits are revealed along the alimentary tract of our sample, each including four intact, unabraded seeds in a regular configuration, as well as another kind of disc-shaped seeds, and no gastroliths are preserved (Figure 4A; Figure 4 - figure supplements 1A). Another frugivore *Bombycilla garrulus* (Bohemian waxwing, Figure 4 - figure supplements 1C, D) was also sampled, and the same situation of the contents is revealed as in *M. comrii*. All the seeds preserved through its alimentary tract including crop, stomach and intestines are intact, and more sparsely located than in the seed-grinders and seed-crackers that we sampled.
2. Seed-cracking parrots: *Conuropsis carolinensis* (Carolina parakeet, Figure 4E), a parrot, is a specialized seed-cracker using beak to de-husk the seeds (Billerman et al., 2020). The alimentary tract of this sample contains a proportionally small bolus of highly fragmented seeds with original shapes impossible to determine, and very few small and sparse stones.
3. Seed-cracking passerines: *Geospiza fuliginosa* (Small ground-finch, Figure 4D, Figure 4 - figure supplements 1E) is half a seed-cracker and half a seed-grinder, and has a diet mostly consisting of small seeds (Billerman et al., 2020). The crop contents of this sample consist of seeds with almost intact configuration, whereas those in the stomach are highly fragmentary along with lots of large gastroliths. We then sampled another seed cracking passerine *Calcarius lapponicus* (Lapland longspur, Figure 4 - figure supplements 1F), and found the same situation of the contents as in *G. fuliginosa*.
4. Seed-grinding granivores: *Ectopistes migratorius* (Passenger pigeon, Figure 4C, Figure 4 - figure supplements 1B), a seed-specialist pigeon, is a seed-grinder that entirely uses gastroliths to crack the seeds (Billerman et al., 2020). Its crop contains numerous, well-defined and intact seeds, whereas seeds are highly fragmented in the stomach, similar to those in *C. carolinensis* and *G. fuliginosa*, together with two large, round gastroliths. Another representative, *Pedionomus torquatus* (Plains-wanderer, Figure 4B, Figure 4 - figure supplements 2A-D) is a general, small-sized seed-grinder. The seeds preserved in the alimentary tract of *P. torquatus* are comparatively more intact than those in other seed specialists such as parrots, pigeons and finches, but many seeds show partial breakages and the gastroliths they contained are much smaller. This indicates that *P. torquatus* might utilize another strategy of abrasion to digest the seeds rather than entirely fragmentation. To test this interpretation, we sampled another seed generalist, *Thinocorus rumicivorus* (Least seedsnipe, Figure 4 - figure supplements 2E, F). The seed remains are in the same condition as in *P. torquatus*

## Data and materials availability

The new specimen reported here (*Jeholornis* STM 3-8) is housed and available for future researchers to check at Shandong Tianyu Museum of Nature, China. The original CT scanning slices and segmented STL files of *Jeholornis* STM 3-8 and involved modern birds, and the alimentary contents of selected modern birds are available in Morphosource (https://www.morphosource.org/projects/0000C1212?locale=en; https://www.morphosource.org/projects/00000C420; https://www.morphosource.org/projects/0000C1080?locale=en). Other data that support this study are available in Figshare (https://figshare.com/s/6a96el338cadd7Of9723), including the rotating videos of the original/reassembled cranial 3D models of *Jeholornis* STM 3-8 and the 3D models of the alimentary contents of selected modern birds, and the landmark data and taxa lists used in GMM analyses. Further information and requests for resources should be

---

**Figure 2 - figure supplements 1.**
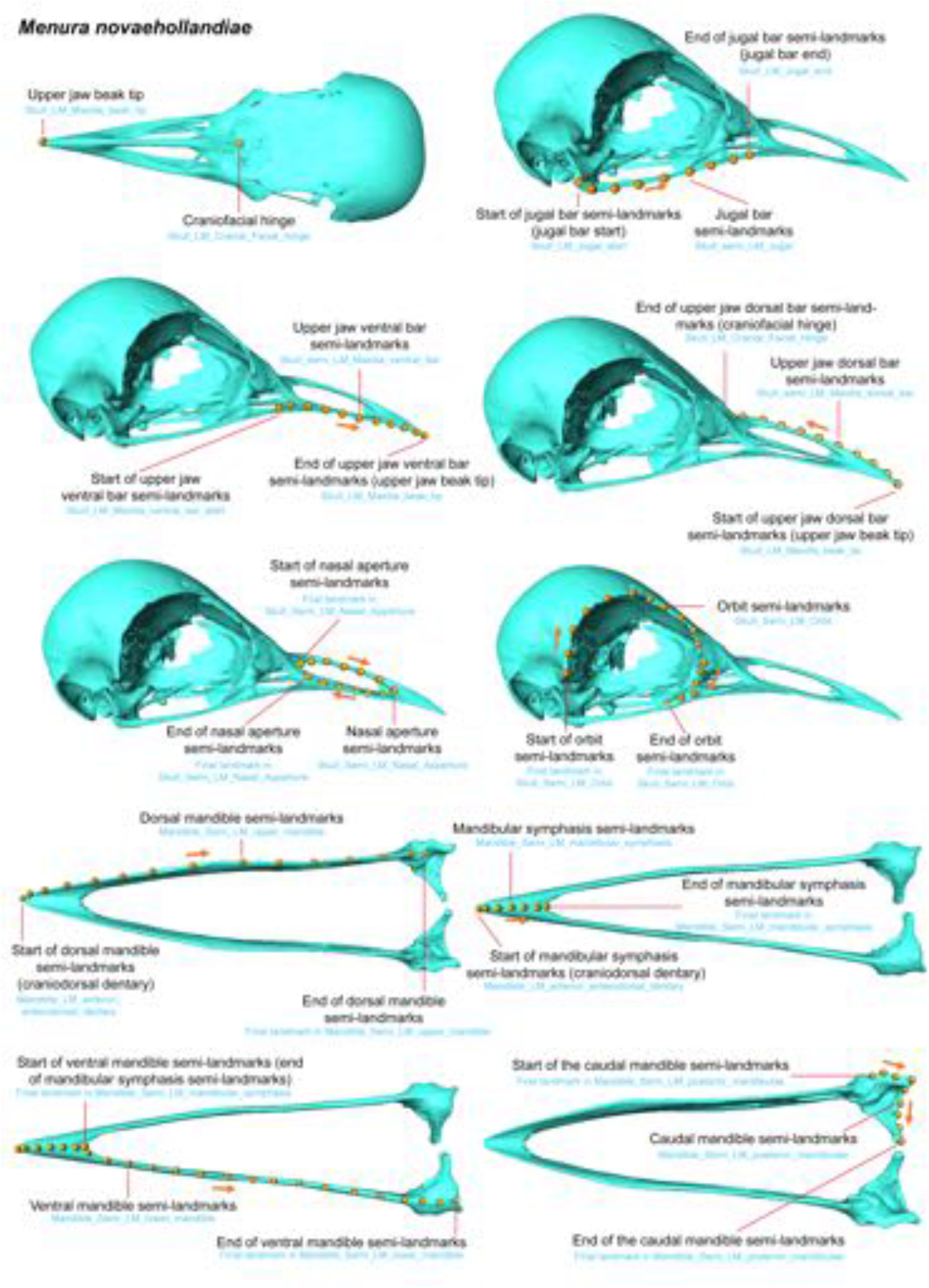
Landmark and semi-landmark locations in *Menura novaehollandiae* as an example of modern taxa used in GMM analyses (following Bjarnason and Benson, 2021). Black labels indicate the description names, and blue labels below indicate the names in R.

**Figure 2 - figure supplements 2.**
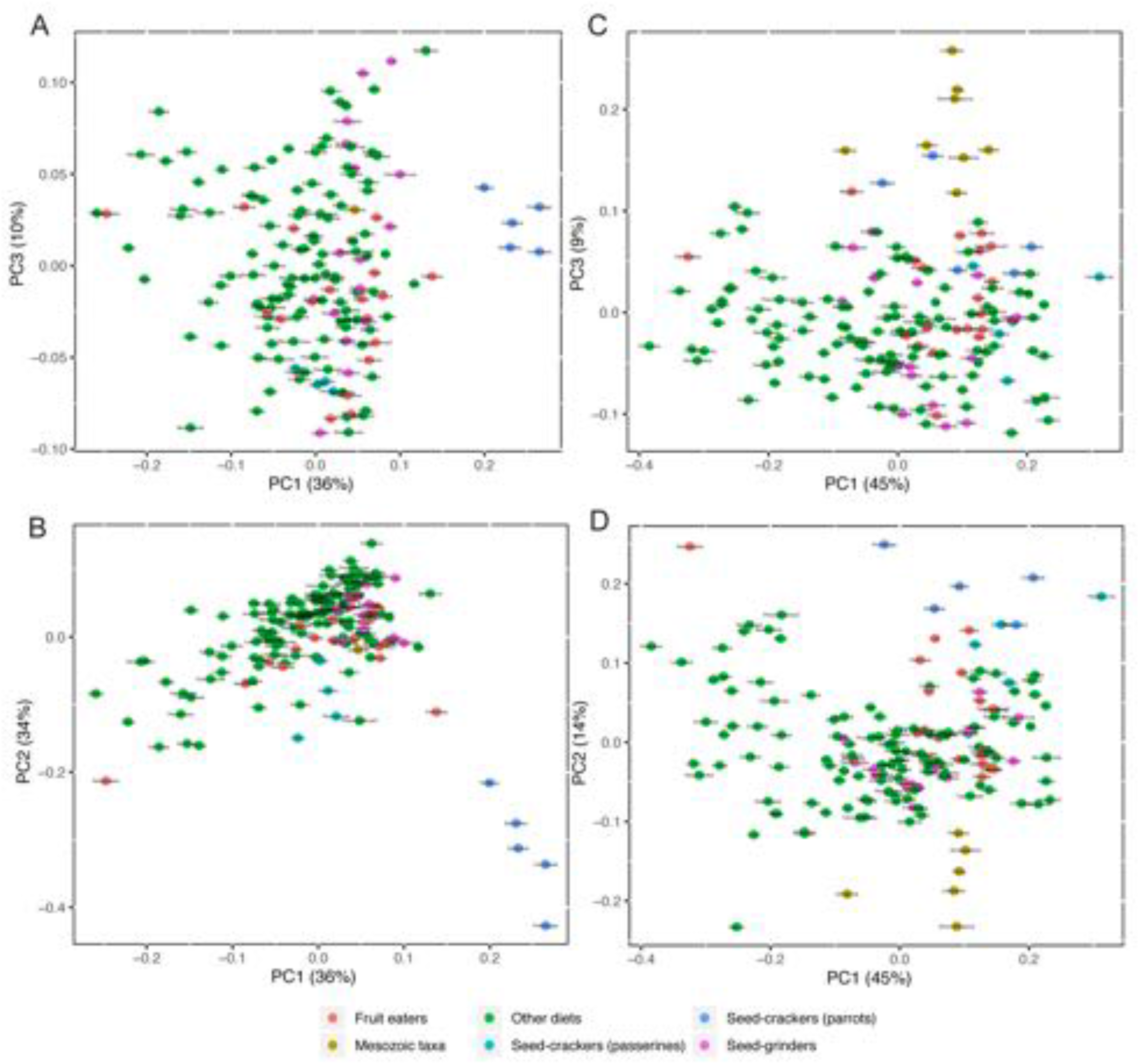
PCA result of 3D mandible (A, B) and 2D skull shape (C, D) with labels of generic names. Some points in green overlap the points in other colours.

**Figure 4 - figure supplements 1.**
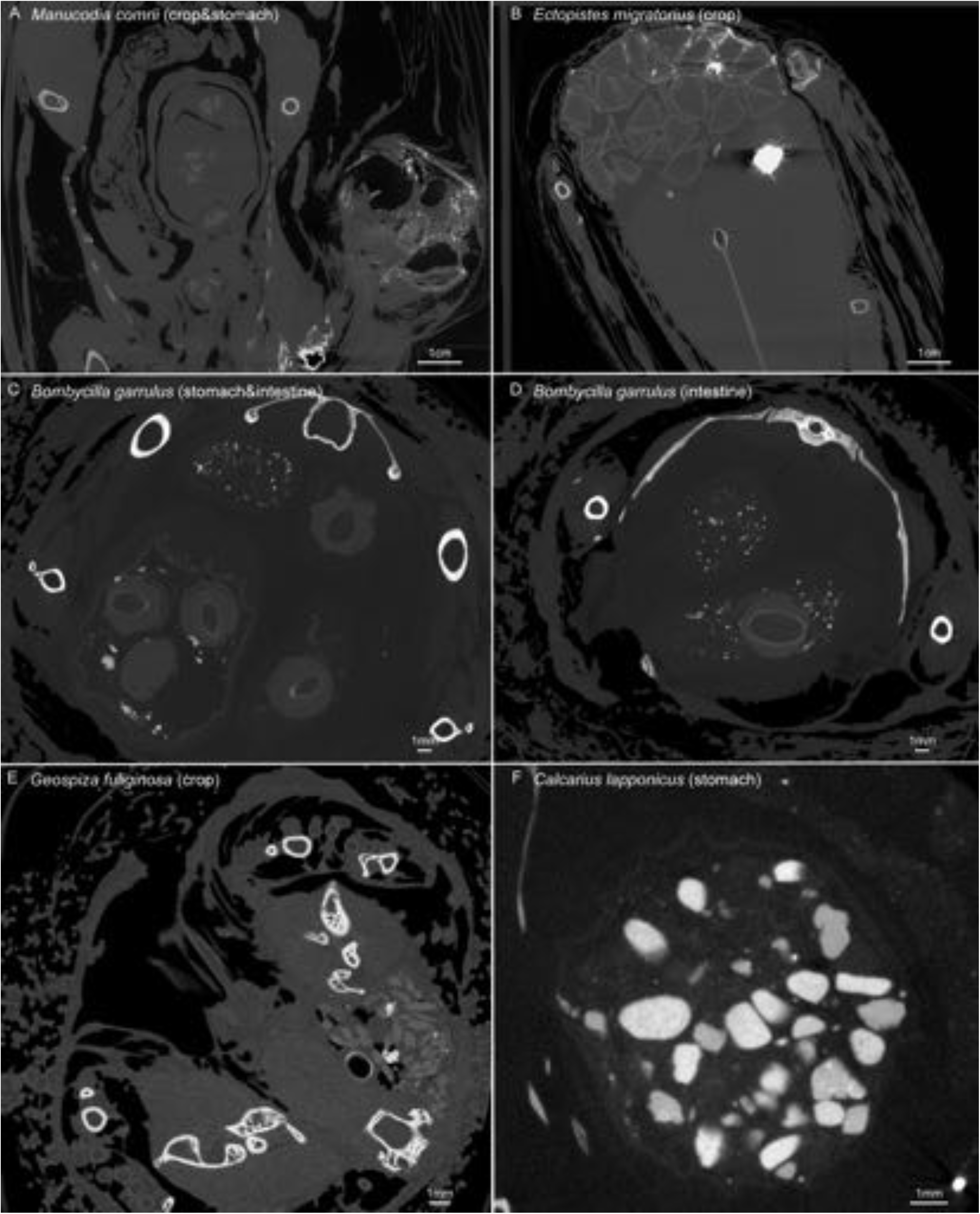
Scanning slices of the alimentary contents in involved modern bird samples.

**Figure 4 - figure supplements 2.**
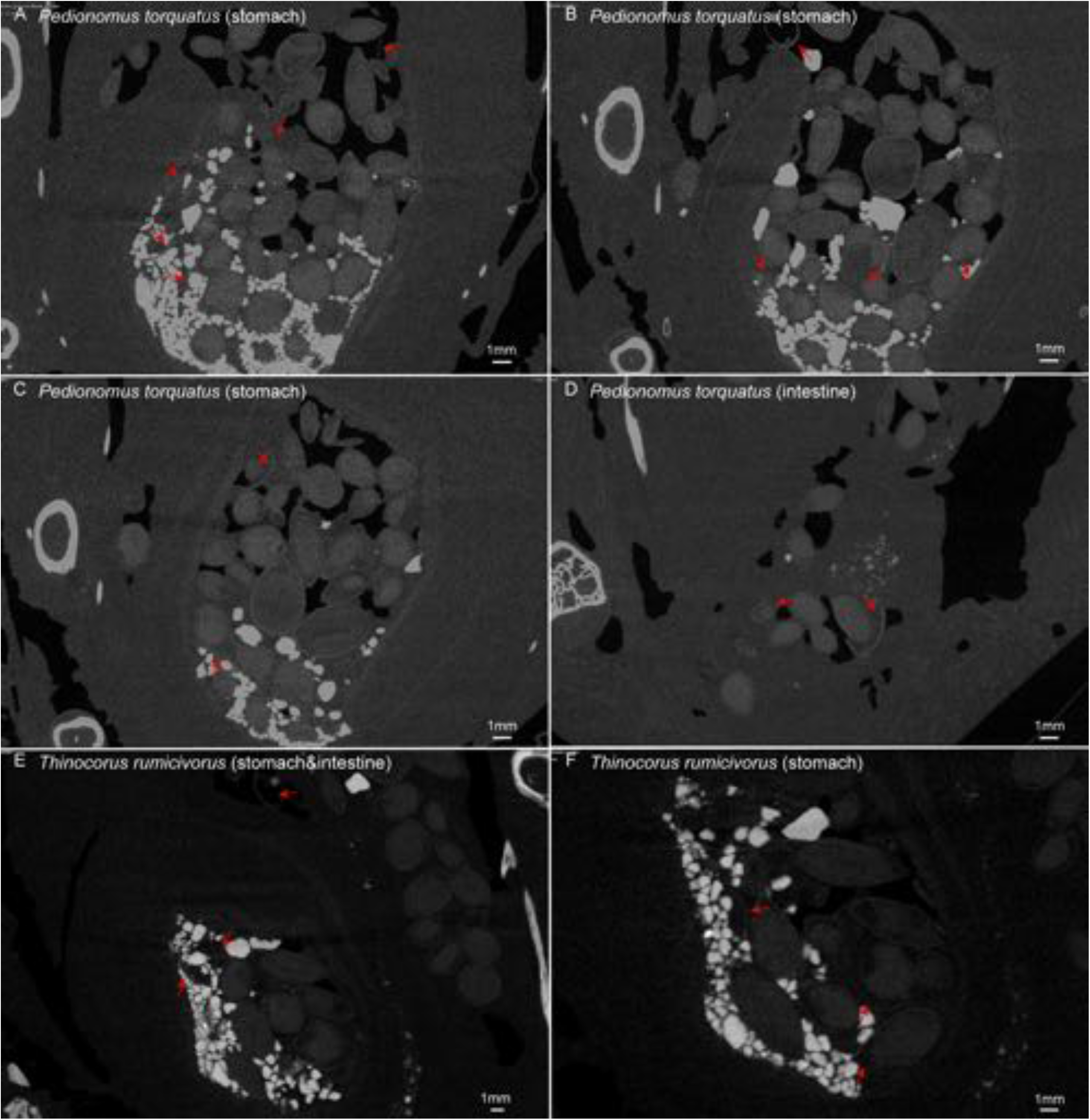
Scanning slices of the alimentary contents in involved modern bird samples. Red arrows indicate the breakages of seeds.

**Figure 2 - Source data 1. Descriptions of cranial and upper jaw landmarks and semi-landmarks** (following Bjarnason and Benson, 2021).

**Figure 2 - Source data 2. Descriptions of mandible landmarks and semi-landmarks** (following Bjarnason and Benson, 2021).

**Figure 4 - Source data 1. Specimens used in the alimentary content analyses.**

